# A rational account of human memory search

**DOI:** 10.1101/326397

**Authors:** Qiong Zhang, John R. Anderson

**Affiliations:** Machine Learning Department, Center for the Neural Basis of Cognition, Carnegie Mellon University, Pittsburgh, PA 15213; Department of Psychology, Center for the Neural Basis of Cognition, Carnegie Mellon University, Pittsburgh, PA 15213

## Abstract

Performing everyday tasks requires the ability to search through and retrieve past memories. A central paradigm to study human memory search is the semantic fluency task, where participants are asked to retrieve as many items as possible from a category (e.g. animals). Observed responses tend to be clustered semantically. To understand when our mind decides to switch from one cluster/patch to the next, recent work has proposed two competing mechanisms. Under the first switching mechanism, people make strategic decision to switch away from a depleted patch based on marginal value theorem, similar to optimal foraging in a spatial environment. The second switching mechanism demonstrates that similar behavior patterns can emerge using a random walk on a semantic network, without necessarily involving strategic switches. In the current work, instead of comparing competing switching mechanisms over observed human data, we propose a rational account of the problem by examining what would be the optimal patch-switching policy under the framework of reinforcement learning. The reinforcement learning agent, a Deep Q-Network (DQN), is built upon the random walk model and allows strategic switches based on features of the local semantic patch. After learning from rewards, the resulted policy of the agent gives rise to a third switching mechanism, which outperforms the previous two switching mechanisms. Our results provide theoretical justification of strategies used in human memory research, and shed light on how an optimal AI agent under realistic human constraints can generate hypothesis about human strategies in the same task.

## 1 Introduction

Performing everyday tasks depend on our ability to retrieve past memories, from finding where the key is, to deciding on a method of transportation in the morning. Very often we need to not only retrieve one specific item but also generate a set of relevant items, from recalling during a hospital visit what were eaten yesterday, to reporting during witness testimony what objects were in the scene. There is rich dynamics in our mental search when we move from recalling one item to recalling the next item. Finding the mechanisms underlying such memory search is important for a better understanding of both how knowledge and information are represented in our memory, and the processes that we use to efficiently navigate through them.

A central paradigm to study human memory search is the semantic fluency task, where participants are asked to retrieve as many items as possible from a category (e.g. animals) in a fixed period. It is observed that responses tend to be clustered semantically (e.g. “cat” follows “dog”) [6]. To understand when our mind decides to switch from one cluster/patch to the next, recent work has proposed two competing mechanisms.

Under the first mechanism, people make strategic decisions to search the semantic space, similar to how animals optimally forage in a patchy spatial environment [6]: one forages locally in one food patch, then switches to a new patch when the resources in the current patch is depleted. It was observed that participants leave a patch in memory search when current rate of finding items is near the average rate for the entire task, consistent with what the marginal value theorem predicts in optimal foraging. The second mechanism, however, demonstrates that similar behavior patterns (i.e. that are consistent with the marginal value theorem) can emerge using a random walk simulation on a semantic network generated by human word-association experiments [1]. This happens irregardless of whether the random walk operates continuously, or when the random walk allows a switch to the start node at a fixed probability at every time step.

To summarize the previous work, both switching mechanisms assume a search process with local/global switch, but differ in their policies of when to switch. The former exploits locally, and make strategic globally switches under the policy of marginal value theory [6]. The latter exploits locally, but has a fixed probability to switch at each time step, which is non-strategic [1]. The two mechanisms give rise to non-distinguishable behavior patterns in human data.

To better compare the two switching mechanisms and propose alternative ones, in this paper, we take a different approach to evaluate different strategies in human memory search. Instead of discussing which model is better supported by human data, we consider the abstract computational problem posed by searching a semantic memory network, and explore what would be an optimal strategy in this task (i.e. which strategy can generate the most items in a semantic fluency task). This approach is based on following principle about human cognition [2][3]:

> Principle of Rationality: Human memory behaves as an optimal solution to the information-retrieval problems facing humans.

A rational analysis of human cognition aim to explain human behavior as an optimal solution to the computational problems posed by our environment ([3]; see also bounded rationality in [16] and ecological rationality in [18]). Examining these computational problems can lead to new models and help us understand better the task at hand. One may argue that human memory does not perform optimally, as we often mis-remember events that are only in the recent past or take a long time to retrieve something that computers can do instantaneously. To clarify, it is therefore important to incorporate the cost and limitations that human cognition faces, and then derive an optimal solution under those considerations [3].

Using a random walk model to simulate a semantic fluency task, we evaluate the task performance under two switching mechanisms. We also propose a third switching mechanism that is consistent with optimal foraging but assumes a policy that is different than the marginal value theorem. It instead bases the exact policy of switching on local patch quality (i.e. statistics around the currently visited node). It is unclear how the local patch quality converts to a switch/not-switch decision that optimizes task performance. Therefore, we will use a reinforcement learning framework with deep Q-learning to obtain the third switching mechanism [11].

The remaining paper is organized as below. We will first describe the random walk model that is used to simulate the semantic fluency task. We will then describe the three switching mechanisms, and how to optimize task performance under each switching mechanism. Then we will evaluate the three switching mechanisms by comparing their relative performances under the simulations. Lastly, we will discuss implications of the results and potential future directions.

## 2 Model

In this section, we will first describe how to use random walks to simulate the semantic fluency task under different switching mechanisms, closely following the procedure used in [1]. Then we will discuss how to optimize task performance under each switching mechanism.

The random walk considered by [1] operated over a semantic network with 5018 words (165 of them are animal names) [12]. This semantic network was obtained from human behavior database in a word-association task, where more than 6,000 participants responded with the first word that came into their mind when cued with another word. The weight of the edge from word A to word B is the proportion of participants that responded A when cued with B. Following closely the procedure described in [1], a random walk over this semantic network searches memory from the start node “animal”. At every single step, it moves from current node to the next node under two options. Under the first option, it transits from the current node *X*_*i*_ to the next node *X*_*j*_ with a probability proportional to the edge weights:

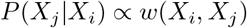

Walking to the neighboring nodes is similar to locally exploiting the resources. Under the second option, it jumps to the node “animal’ and continues to a next node from there. Opportunity to switch away from the local patch allows for global exploration. The three different switching mechanisms we will compare specify the exact policy for local/global switch at each time step.

The semantic fluency task requires responses of non-repeated animal names. The random walk generates such responses every time it first visits an animal name. Therefore, the time it takes between two adjacent animal names generated, i.e. inter-response times (i.e. IRT), can be obtained as:

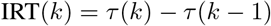

where *τ*(*k*) is the first visiting time of *k* th animal name generated. For example, if the first five visited nodes of the random walk are *X*_0_ = “dog″, *X*_1_ = “house″, *X*_2_ = “cat″, *X*_3_ = “dog″, *X*_4_ = “mouse″. In this case, *k* = 1 refers to “dog” with *τ*(1) = 1; *k* = 2 refers to “cat” with *τ*(2) = 3, since it skips a node “house” that is non-animal; *k* = 3 refers to the third valid response “mouse” with *τ*(3) = 5, since it skips a node “dog” which has already been visited. Therefore, we can calculate the time between generating the third response “mouse” from the preceding response “cat”

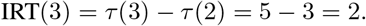

In this way, we can simulate the semantic fluency task and give behavior results (i.e. a sequence of generated animal names and its inter-response times) similar to those in human experiments.

### 2.1 M1: Switch under marginal value theorem

The first switching mechanism (M1) is based on the marginal value theorem, which is an important model that characterizes optimal foraging behavior in the literature of animal foraging [4]. It is based on the assumption that resources are monotonically depleted during foraging. Under this environment, animals seek to maximize the gain per unit time R during foraging, defined as below:

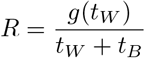

where *g*(*t*_*W*_) is the cumulative gain within a patch, *t*_*W*_ is the time spent within each patch, and *t*_*B*_ is the time spent traveling between patches.

To optimize *R*, marginal value theorem describes that the optimal foraging policy is to leave a patch when the instantaneous rate (i.e. marginal value) of gain is equal to the long-term average intake under the current environment. When it transfers to memory search, marginal value theorem predicts that individuals should leave the current patch and switch to another when the gain falls below the expected benefits of searching elsewhere in memory (see more details in [6]). Given fixed reward per retrieved item, the current switching mechanism describes that one should leave a patch once the instantaneous time cost (i.e. time cost of the just-retrieved item) is equal to the average time cost under the environment.

To optimize the task performance under this switching mechanism, we will explore a range of time thresholds, and select the one that demonstrates the best task performance. This is to relax the original marginal value theorem to one that relies on using a fixed-time threshold, since we do not have knowledge of the exact average cost prior to the completion of the task.

### 2.2 M2: Switch at a fixed probability

The second switching mechanism (M2) describes a scenario where local/global switch takes place in a random and non-strategic way [1]. At each time step, there is a fixed probability *ρ* that the random walk jumps back to the start node “animal”; the other times under probability (1 − *ρ*), it transits from the current node to the next node, with a probability proportional to the edge weights. To optimize the task performance under this switching mechanism, we will explore a range of values *ρ*, and select the one that demonstrates the best task performance.

### 2.3 M3: Switch based on local patch quality

We propose a third switching mechanism (M3) based on the local patch quality, described by local statistics around the currently visited node in the random walk. The intuition is straight-forward: one switches to a new patch whenever the resources in the current patch is depleted. This switching mechanism borrows ideas from the observations in other human foraging tasks outside the memory domain: under a virtual environment where human participants forage for food, their behavior is sensitive to the absence and presence of the resources locally [9] [7]. Deciding to switch based on local patch quality is also consistent with the observations in [6], where there is lowest residual proximity (defined as the average similarity to non-visited neighbors) prior to a local/global switch.

To optimize the task performance under this switching mechanism, we need to obtain a random walk agent that best performs the task while utilizing the information from local patch quality. However, unlike M1 and M2 with specified parameters, it is unclear how the information of various local statistics of the currently visited note converts to a switch/not-switch decision. Therefore, we will use a reinforcement learning framework to learn a best model under the switching mechanism M3. We consider that a random walk agent interacts with the environment in a sequence of actions, observations and rewards. At each time step, the agent selects an action *a*_*t*_ from a binary set *A* = {0,1}, which represents whether to switch or not. The entire semantic graph that the random walk operates on is not visible to the agent. Instead, the agent only has information of the immediate neighbors of the node that is currently visited, assuming minimally about what information can be accessed during human memory search. Figure 1a plots the histogram of the number of neighbors of every node in the network. It can be seen that the semantic network based on free association norms are very sparse. Figure 1b plots the histogram of the sum of edge weight of the top five neighbors, which consisted of a majority of the total weight (i.e. 1) in most of the nodes examined. Therefore, we assume that the agent only need to access local patch information associated with a few nearest neighbors (i.e. top five). There can be three types of information associated with each neighbor node, including 1) the edge weight, 2) whether it is a non-visited animal node, and 3) how many times it has already been visited. This gives rise to state s consisted of a vector of 15 values. One simulated trial is considered as one episode, which terminates when the time is up. We consider a range of time limits *T*_0_ = {500,1000, 2000}, which leads to performance level of the agent at the same scale as the human experiments [6]. At each time step *t*, the agent receives a reward *r*_*t*_ = {0,1}, which represents whether a new animal name is successfully generated. This formalism leads to a finite Markov decision process, which we can apply standard reinforcement learning methods to.

The goal of the agent is to find the switching mechanism that selects switch/not-switch in a fashion that maximizes cumulative future reward (discounted by a factor of γ per time step). This rule can be obtained from the optimal action-value function:

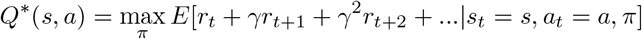

which is the maximum sum of rewards *r*_*t*_ achievable by a behavior policy π = *P*(*a*|*s*), after making an observation *s* and taking an action *a*. *Q*\*(*s*, *a*) obeys the Bellman equation:

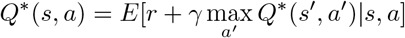

where *s′* at the next time step is known for all possible actions *a′*. Using Bellman equation as an iterative update, i.e. *Q*_*i*+1_(*s*, *a*) = *E*[*r* + γmax_*a′*_ *Q*_*i*_(*s′*, *a′*)|*s*, *a*], it will converge to optimal action-value function *Q*_*i*_ → *Q*\* as *i* → ∞ [17]. To allow the representation to emerge flexibly from experiences, we use a deep neural network to approximate the action-value function *Q*(*s*, *a*; θ) ≈ *Q*\* (*s*, *a*). This results in a Deep Q-Network (DQN) with weights represented by *θ* [11]. It is trained by adjusting the parameters *θ*_*i*_ at iteration i to reduce the mean-squared error in the bellman equation. This leads to a loss function updated at each iteration *i*:

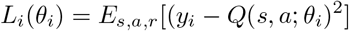

where the target for iteration *i* is *y*_*i*_ = *E*_*s′*_ [*r* + γ max_*a′*_ *Q*(*s′*, *a′*; *θ*_*i*−1_)]. We then use the Q-learning algorithm to update weights by optimizing the loss function with stochastic gradient descent [20]. The behavior distribution during training uses a e-greedy strategy, with follows greedy strategy with probability (1−*∊*) and selects a random action with probability *∊*. To model the switching mechanism as a simple heuristics like that in M1 and M2, we adopt a very simple neural network architecture, which is consisted of only two hidden layers with ten and five hidden units respectively.

To summarize, the obtained DQN describes the third switching mechanism. At each time step, the action to take is the one with largest Q-network value under the current *s* (local patch quality).

**Figure 1:**
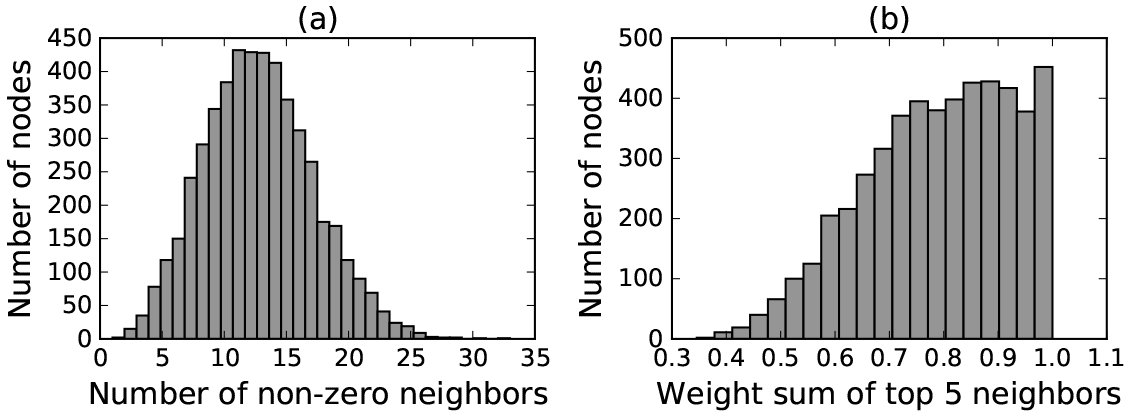
Histogram of the number of neighbors of every node in the semantic network (a). Histogram of the sum of edge weight of the top five neighbors (b)

## 3 Results

There are two parts for the results section. The first part is the behavior level results, where we compare the three switching mechanisms based on the generated animal names and inter-retrieval times. The second part is the rational analysis, where we evaluate the three switching mechanisms based on their performances in the semantic fluency task.

### 3.1 Three switching mechanisms give similar markers at the behavior level

Three rows in Figure 2 demonstrate the behavior results under three switching mechanisms M1, M2, and M3 respectively. The first column shows that all three switching mechanisms give inter-response times (IRTs) consistent with the marginal value theorem. Switches were identified using pre-defined categories from [19]. Marginal value theorem would predict at the behavior level that, switches take place when the instantaneous IRT exceeds the average time cost. This is consistent for all three switching mechanisms, where we observe IRT equals or exceeds the average cost right before the switch (i.e. “−1” relative to switch). It is then followed by a larger time cost right after the switch, and a decreased time cost for the next items in the new patch. The second column shows the individual episodes of the simulations (analogous to simulated trials in the human experiment), where the average IRT and last item IRT prior to switch lie at the diagonal line, also consistent with the marginal value theorem. The third column shows that for individual episodes of simulations, those with smaller difference between average IRT and last item IRT gives better performance on that episode, with correlation coefficient of −0.21 and −0.17 in M1 and M2, but with an exception in M3 with −0.02. The lack of effect in M3 is a result of our optimizing performance on each episode. One would need to simulate individual differences in performance by introducing deviation from the switching mechanism specified by M3. To summarize, carrying out the same analysis as [6] and [1], we observe similar behavior evidence across all three switching mechanisms. Though only M1 is generated based on marginal value theorem, all three switching mechanisms give results consistent with what marginal value theorem would predict. This suggests that we need other methods to compare three switching mechanisms, which is what we will do in the next section with a rational analysis.

### 3.2 Switching mechanism M3 based on local patch quality outperforms M1 and M2

Figure 3a shows that the reinforcement learning agent under M3 performs better than M1 using marginal value theorem. This is the case with different settings of time limits, 500, 1000, and 2000. The threshold (T) used in marginal value theorem has minimal effect on the overall performance (number of items recalled).

Figure 3b shows that the reinforcement learning agent under M3 performs better than M2 under probabilistic switch. This is the case with different settings of time limits, 500, 1000, and 2000. With increasing switching probability, the performance first increases and then decreases, indicating that there is an optimal switching probability where it achieves the best exploration/exploitation trade-off.

Figure 3c shows the training curve tracking the RL agent’s task performance when the time limit of the task is 2000. After around 1000 episodes, the RL agent under M3 achieves performance that is better than both M1 and M2.

**Figure 2:**
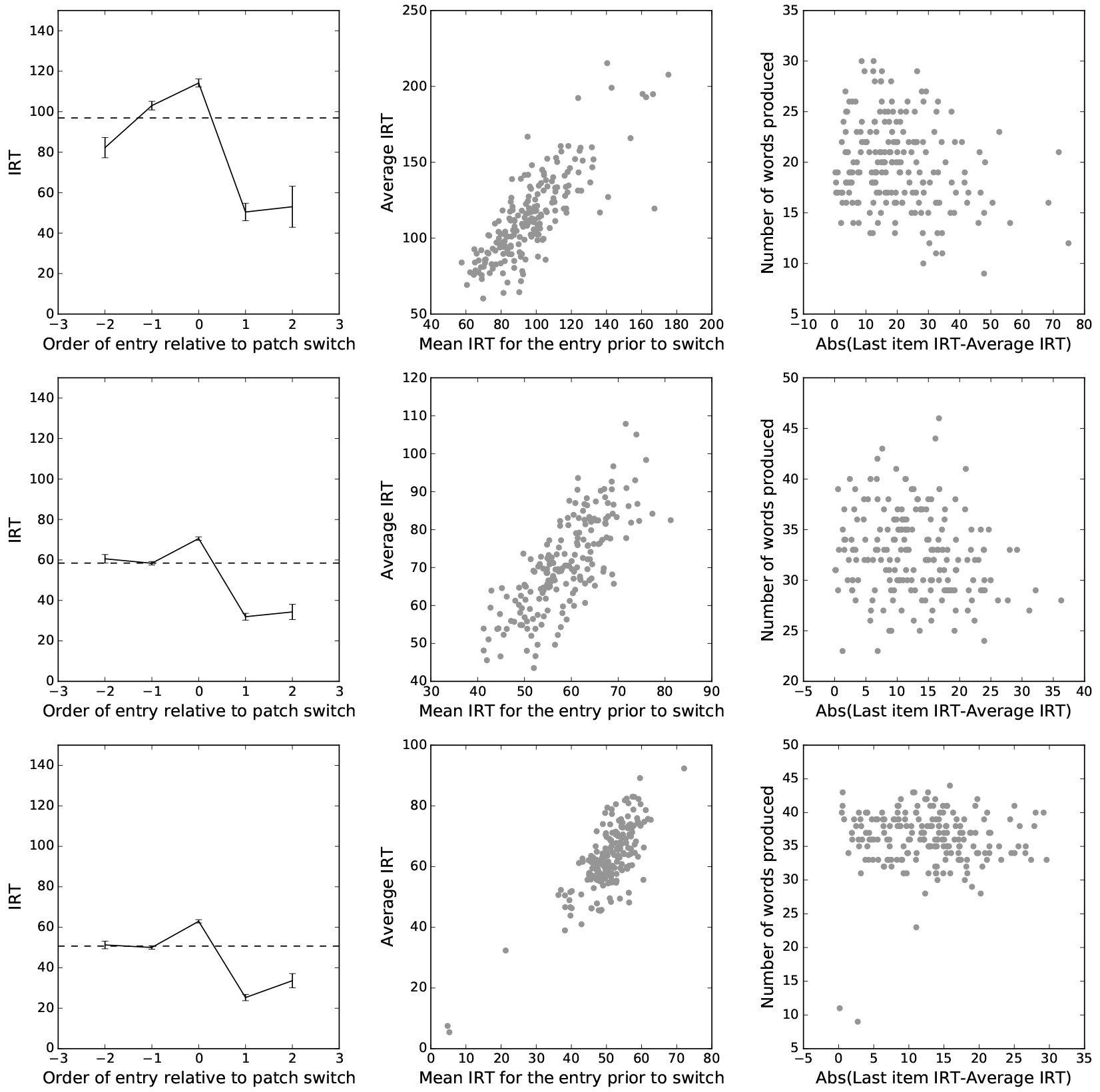
Three three rows correspond to behavior results under M1, M2 and M3, each with 200 simulations. First column plots averaged IRTs around switches, and the average IRT over the entire task (dotted line). Second column plots for each simulated episode the average IRT versus the last item IRT prior to switch. Third column plots the episode performance versus the absolute difference between the average IRT and the last item IRT prior to switch.

**Figure 3:**
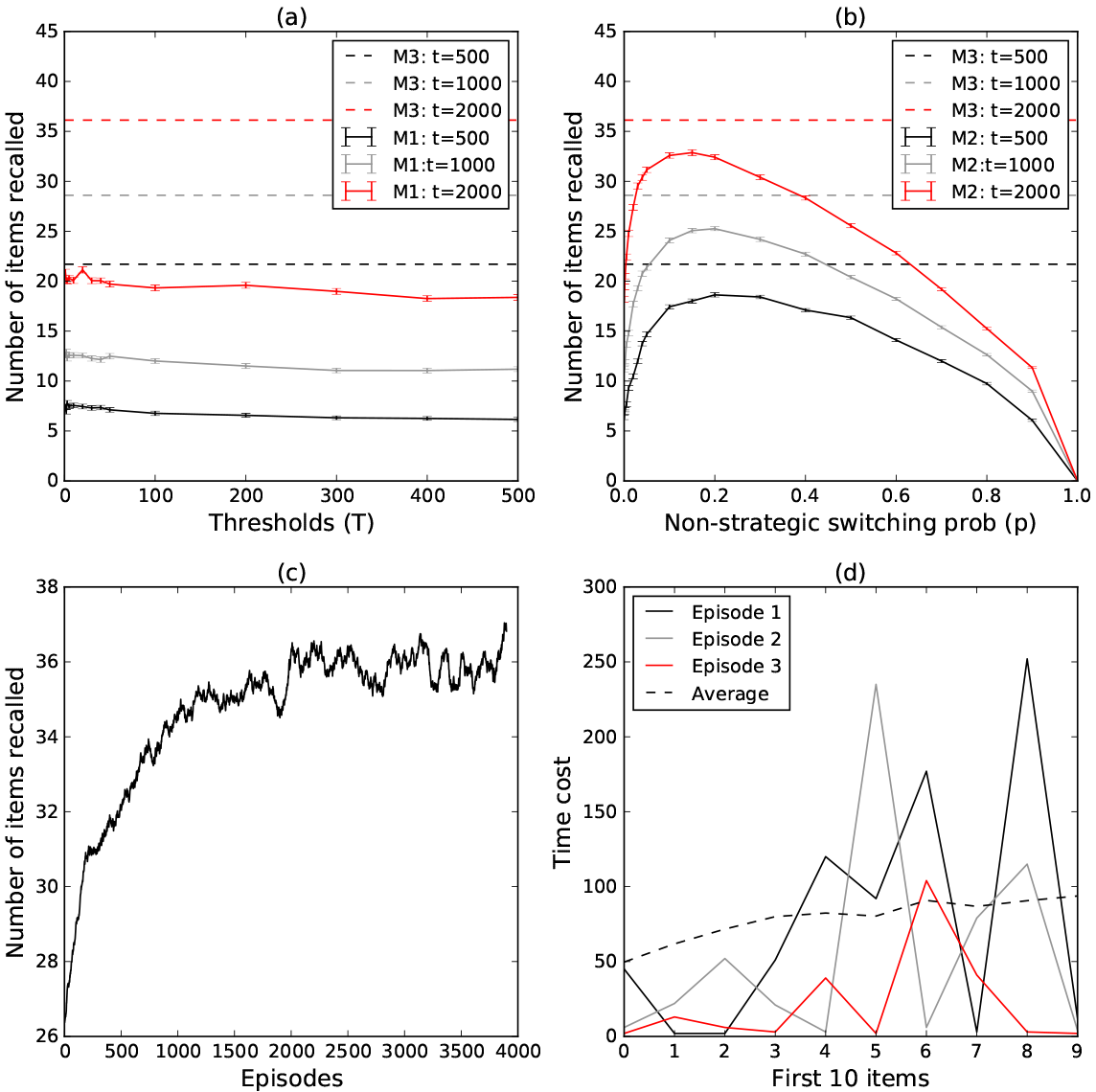
Comparing performance of M3 with that of M1, under different time limits, 500, 1000, and 2000 (a).Comparing performance of M3 with that of M2, under different time limits, 500, 1000, and 2000 (b). Training curve tracking performance of M3 when *t* = 2000, averaged over a window of 100 episodes (c). IRTs for the first ten items for three representative episodes or averaged across all episodes (d). Hyper-parameters used in the DQN under M3: learning rate 0.01, number of episodes 2000, γ = 0.95, *∊*_max_ = 1, *∊*_min_ = 0.0001, *∊*_decay_ = 0.999.

Figure 3d plot the IRTs for the first ten items for three representative episodes. They are compared with the IRTs for the first ten items averaged across all episodes. One can observe that time cost is only monotonically increasing in the averaged IRTs but not for IRTs within individual episodes.

## 4 Discussion

Despite evidence of existence of local/global switches [6], an exact decision rule of when a switch takes place is unclear. Recent work has proposed two competing accounts to characterize the memory search patterns in the semantic fluency task [6][1], which converts to two distinct switching mechanisms. Our contribution is two folds in this study. First, we propose a third switching mechanism in memory search based on the local patch quality. Second, we carry out a rational account of what would be an optimal strategy in the memory search. Comparing the task performance of three switching mechanisms in the semantic fluency task, our proposed switching mechanism based on the information of local patch quality outperforms the other two. Based on the rational analysis [2][3], human cognition optimally solves the problems that it faces; therefore, the mechanism that has an advantage in achieving better task performance would be closer to what human memory does. Our results provide theoretical justification of strategies used in human memory research, and shed light on how an optimal AI agent under realistic human constraints can generate hypothesis about human strategies in the same task. Given that the current study is purely theoretical, below we will discuss the plausibility of representation and learning mechanism underlying the proposed switching mechanism, and the potential reasons why the other two switching mechanisms do not perform as well. Lastly, we will discuss limitations and future directions of the current work.

### 4.1 Plausibility of the proposed switching mechanism in human memory

One may wonder if the human mind is capable of obtaining the representation like that in the DQN in the current task. We will discuss in turn the three components about the DQN we used to obtain the switching mechanism M3. The first component is the temporal differencing that learning Q-network is based on. In the literature of human reinforcement learning, there has been neural evidence of temporal differencing identified, suggesting that human are capable of using temporal differencing to learn from sparse future rewards [14][13].

The second component is the neural network that represents Q-function. It is possible for human to adopt such a representation as it assumes very minimally about the computations it does, with only two hidden layers and fifteen hidden units in total. The simplicity of the neural network architecture also ensures that the learned switching mechanism is a simple heuristics like that in M1 and M2. We use a neural network instead of other specified structures, because this allows the optimal underlying representation to emerge from experiences and learning. Similar approaches are used in connectionist models of semantic learning and language acquisition [5][10][15].

The third component is the input data, including both the amount of training experiences and the dimension of the neural network input. We assume very minimally about the amount of information one can access at a time in the current memory, namely, only 15 units representing the local statistics of the currently visited node. However, we do assume that human have been exposed to a large amount of experiences of similar tasks, as seen from Figure 3d. This assumption comes from the rational analysis [2][3], which describes that human cognition optimally adapts to the environment and the tasks it faces. Despite that the specific task of retrieving animal names is not a natural task that we are usually exposed to, there are a lot of similar memory search tasks that we do get exposed to throughout our daily life and development. The Q-network that is eventually obtained through this exposure can be a simple heuristics used in memory search in general.

### 4.2 A comparison with non-strategic switching and marginal value theorem

Non-strategic switching does not perform as well as the switching based on local patch quality. Rational analysis describes that human optimally solves the problems it faces. Therefore, intuitively, if there is a chance of improving the performance through strategic switches (like M3 that uses local patch quality), it is very unlikely that human memory will keep it in a non-strategic way (like M1 that switches at a fixed probability).

Marginal value theorem does not perform well in the simulations, neither is the performance sensitive to the threshold (*T*) used in the switching mechanism. This is potentially contributed by a violation of the assumptions underlying the marginal value theorem. Marginal value theorem assumes that resources are monotonically depleted during foraging [4]. Inter-item retrieval times, while averaged across trials, do give monotonically increasing trend (Figure 3d]). However, there is a lot of variability from one item to the next item of their retrieval times on single-trial basis. Therefore, using marginal value theorem is not a good decision rule on single-trial basis.

### 4.3 Limitations and future directions

When committing to a particular algorithm (i.e. switching mechanism) for memory search, we also make assumptions on the representation the algorithm operates on. Hills et al. focused on a spatial representation with BEAGLE similarity [6][8], which is different than the semantic network used in [1] and the current study. Future work will include extending the current framework of rational analysis to the spatial representation under BEAGLE similarity, and examine if the switching mechanism based on local patch quality is still the most optimal rule under the semantic memory search. Current analysis is a theoretical analysis that is used to generate alternative cognitive mechanisms. Therefore, the second future direction includes testing the proposed cognitive mechanism over observed human data.

